# Limits to the adaptation of herbivorous spider mites to metal accumulation in homogeneous and heterogeneous environments

**DOI:** 10.1101/2023.03.15.532545

**Authors:** Diogo P. Godinho, Inês Fragata, Agnieszka Majer, Leonor R. Rodrigues, Sara Magalhães

## Abstract

Metal accumulation is used by some plants as a defence against herbivores. Yet, herbivores may adapt to these defences, becoming less susceptible. Moreover, ecosystems often contain plants that do and do not accumulate metals, and such heterogeneity may affect herbivore adaptation. Surprisingly, few studies have tested this. Tomato plants accumulate cadmium, affecting the performance of the herbivorous spider mite *Tetranychus evansi*. Here, we performed experimental evolution to test whether these mites adapt to plants with high cadmium concentrations, in homogeneous (plants with cadmium) or heterogeneous (plants with and without cadmium) environments. We measured fecundity, hatching rate and the number of adult offspring after 12 and 33 generations and habitat choice after 14 and 51 generations, detecting no trait change, which implies absence of adaptation. We then tested whether this absence of adaptation was due to a lack of genetic variation in the traits measured and, indeed, additive genetic variance was low for the measured traits. Possibly, we did not measure the traits that contributed to population persistence on plants with cadmium. Interestingly, despite no signs of adaptation we observed a decrease in fecundity and number of adult offspring produced in cadmium-free plants, in the populations evolving in environments with cadmium. Being this the case, evolving in environments with cadmium may reduce the growth rate of spider mite populations on non-accumulating plants as well. Nevertheless, adaptation to metal accumulation may occur via herbivore traits not commonly measured, which calls for broader studies on this topic.

## Introduction

Plant herbivore interactions have long been used as examples of evolutionary arms races, being at the very birth of the term in the classical paper where it was used for the first time (Ehrlich & Raven, 1964). Indeed, plants have evolved a myriad of defences to reduce herbivory, such as toxic compounds or physical structures that hinder herbivore performance (Walling, 2000). Herbivores, in turn, adapt to such defences, by avoiding highly defended plants, improving detoxification or manipulating the expression of some defences (Musser *et al*., 2002; Després *et al*., 2007; Perkins *et al*., 2013).

Besides organic defences, plants may also rely on metals they extract from the soils and accumulate on their leaves (i.e. hyperaccumulation (Baker & Brooks, 1989)) as a defence against herbivores (Martens & Boyd, 1994). Indeed, laboratory studies have shown that metal accumulation by plants has negative effects on the performance of herbivores (Martens & Boyd, 1994; Boyd & Moar, 1999; Jhee *et al*., 2005; Kazemi-Dinan *et al*., 2014) and field surveys show that such plants suffer less herbivory than neighbouring non-accumulating plants (Martens & Boyd, 2002; Noret *et al*., 2007; Galeas *et al*., 2008).

The high mortality imposed by metal hyperaccumulation poses a strong selection pressure upon herbivores to resist or tolerate such metal-based defences. Yet, unlike the vast evidence documenting responses of herbivores to organic plant defences, clear demonstrations of herbivore adaptation to metal defences are limited. It is known that herbivore taxa vary in their susceptibility to metal accumulation (Jhee *et al*., 2005; Konopka *et al*., 2013) and that such variation can be also found within the same herbivore species, but the evidence is as yet scarce (Freeman *et al*., 2006; Xu *et al*., 2021; Godinho *et al*., 2022a). Moreover, most studies do not establish a link between susceptibility and previous exposure to metal defences. Finally, the mechanisms behind herbivore adaptation to metal accumulation are poorly understood. The exception is for one population of the moth *Plutella xylostella* collected on the selenium accumulating plant *Stanleya pinnata* (Freeman *et al*., 2006). This population differs from another population of the same species, naïve to selenium, by being able to methylate selenium, which makes this metalloid less toxic (Freeman *et al*., 2006).

Metal contamination in the soils typically occurs either in gradients, in the case of anthropogenic origin, such as mines, or in patchy landscapes such as serpentine soils (Whittaker, 1954; Notten *et al*., 2005). Moreover, for the same metal concentration in the soil, there is intraspecific and interspecific variation in metal accumulation by plants (Macnair, 2002; Assunção *et al*., 2003). Therefore, hyperaccumulation is highly heterogenous even within small spatial scales, which may affect herbivore adaptation. Indeed, whereas homogeneous environments typically select for specialists, whether a generalist or specialists will evolve in heterogenous environments depends on the strength of the trade-off between adaptation to each environment, on the type of population regulation and on gene flow between environments (Kassen, 2002). Moreover, if differences in productivity between environments is high, organisms may not adapt to the poorest, sink environment (Bisschop et al., 2019; Laska et al., 2021).

Spider mites are herbivorous pests of many crops worldwide and model organisms for the study of plant-herbivore interactions (Grbić *et al*., 2011). Indeed, the effect of plant defences on spider mite life history traits as well as the behavioural and mechanistic responses of spider mites to such defences are well described (Magalhães *et al*., 2007; Kant *et al*., 2015; Godinho *et al*., 2016; Rioja *et al*., 2017). Due to their short life cycle and high growth rates, spider mites are amenable to experimental evolution studies, a methodology that has been used to study, among other topics, their adaptation to novel host plants (Sousa *et al*., 2019). Furthermore, metal accumulation by different plant species has a negative effect on the growth rate of spider mites (Jhee *et al*., 2005; Quinn *et al*., 2010; Godinho *et al*., 2022b).

*Tetranychus evansi* is an invasive crop pest in the Mediterranean basin, being a specialist of solanaceous plants and having the remarkable ability to suppress defences in tomato plants (*Solanum lycopersicum*), improving herbivore performance on such plants (Sarmento *et al*., 2011; Godinho *et al*., 2016). However, despite suppression continuing to occur in the presence of metals, this species is negatively affected by cadmium accumulation in tomato plants (Godinho *et al*., 2018) and it is unknown whether the adaptation of this herbivorous spider mite to cadmium accumulation in tomato plants is possible. Here, we tested the adaptation and costs thereof, of *T. evansi* evolving in an environment with cadmium accumulating tomato plants and in a heterogeneous environment, composed of plants with and without cadmium. Additionally, in order to characterize the genetic potential for adaptation to cadmium accumulation, we estimated the amount of genetic variance present in the control population in response to cadmium.

## Materials and methods

### Plants

Tomato plants (*Solanum lycopersicum*, var Moneymaker) were kept in a climatic chamber (photoperiod of 16:8h, 25:20°C, day:night). They were sown in germination soil (SIRO, Portugal) and, two weeks later, transplanted to a mixture of 1:4 vermiculite, gardening soil (SIRO, Portugal). Subsequently, plants grew 3 more weeks, being watered twice a week with 100 ml of distilled water or 100 ml of a solution of 2mM cadmium chloride, and one additional time with tap water to compensate for micronutrients deficiencies (Godinho *et al*., 2018)

### Ancestral population

In this study, we used an outbred population of *T. evansi*, created by merging 4 natural populations via one-on-one controlled crosses (Godinho *et al*., 2020). This population was kept at high numbers (> 1000 adult females) on a cage containing two entire tomato plants, at 25 ºC ± 1 and 16:8 hours of light:darkness. Every other week, one tomato plant was replaced with a fresh one to keep maintain *ad libitum* food provisioning.

### Selection regimes

From this outbred population, three selection regimes were created, each consisting of four detached leaves of tomato plants collected from different plants and grown in soils with or without cadmium. Because tomato plants accumulate cadmium on their leaves (Godinho *et al*., 2018), this resulted in leaves with or without cadmium. The three selection regimes were: a control environment with cadmium-free leaves, a homogenous environment with high cadmium (i.e., with leaves from plants grown with a 2mM cadmium chloride supply) and one heterogenous environment, in which half of the leaves contained cadmium and the other half did not. To compensate for nutritional and defensive differences pertaining from leaf age, each environment was composed of leaves of different ages (2^nd^ to 5^th^ counting from the cotyledons). In the heterogeneous environment, we used the following combinations: a) 2^nd^ and 4^th^ leaves without cadmium and 3^rd^ and 5^th^ leaves with cadmium or b) 3^rd^ and 5^th^ leaves without cadmium and 2^nd^ and 4^th^ leaves with cadmium. The combinations were alternated among replicates of the same selection regime and every other generation.

For each selection regime, five independent replicate populations were founded with 220 mated females from the outbred population and maintained in discrete generations by transferring the same number of females every 14 days to a box containing fresh leaves. The remaining individuals from each population were kept in separate boxes (designated t-1) for one more generation with relaxed selection (i.e., on leaves without cadmium), so that they could be used as a backup in case there was any sudden drop in mite number that prevented the maintenance of a selection regime. Thus, at the time that mites would have to be transferred to a new box to form the next generation (t), if there were less the 220 mated females in the population, the remaining individuals would be transferred from the equivalent (t-1) box (Godinho *et al*., 2020). If still not enough individuals were available to complete a transfer, the remaining were collected from the ancestral outbred population. This allowed us to maintain the same population size across replicates and selection regimes. Given that the number of mated females transferred at a given generation from the (t), the (t-1) and the ancestral populations varied among replicates, for each experimental population the effective number of generations of selection was determined for each timepoint at which the different assays were performed (corresponding to 12, 14, 33 and 51 discrete generations, assuming no backup was necessary) according to Godinho *et al*. 2020 (Fig. S1). The effective number of generations of selection was then incorporated as a covariate in all models (Table S1).

The maintenance and transfer of the experimental populations was done in 5 blocks, each consisting of one replicate population of each selection regime.

All experimental populations were kept in a climatic chamber at 26 ± 1 ºC, with a 16:8 hours of light:darkness photoperiod. Before each experiment, individuals from each replicate experimental population were collected and kept in a cadmium-free common garden environment for 2 generations to homogenize maternal and environmental effects across individuals from different selection regimes. After this period, mated females were collected, circa 2 days after emerging from the last quiescent stage and used in the experiments, except for the “quantification of genetic variation” experiment, in which quiescent females were collected to ensure virginity. All common gardens and experiments were maintained in a climatic chamber at 25 ± 1 ºC, with a 16:8 hours of light:darkness photoperiod.

### Spider mite performance

Mated females were isolated on leaf discs (Ø 18 mm) made from plants supplied or not with 2mM cadmium chloride. These leaf discs were kept on top of soaked cotton wool in petri dishes. 48 hours later, females were killed, and the number of eggs was counted. One week after instalment of the mated females, hatching rate was registered and, 14 days after instalment, the number of adult offspring was counted. This experiment was performed after 12 and 33 discrete generations of experimental evolution.

### Habitat choice

Mated females were placed on the centre of parafilm strips (1×3 cm) connecting two leaf discs (Ø 16 mm) placed on soaked cotton wool, one made from a tomato plant supplied with cadmium (2 mM) and the other from a plant without cadmium. On each trial, leaf discs were made from leaves of the same age (either the 3^rd^ or 4^th^ counting from the cotyledons) and the age and the position of the disc with cadmium was randomized across trials. These trials were performed at discrete generations 14 and 51. At generation 14, 24 hours after the instalment, the position of the female and the number of eggs laid on each disc were registered. At generation 51, the position of the female was registered after 24 and 48 hours, and the number of eggs laid was only counted after this period.

### Genetic variation

Genetic variation in the number of eggs and total offspring in the presence of high cadmium concentrations was quantified by performing a full-sib half-sib experiment (following the approach described in Magalhães *et al*., 2007) using one of the replicate populations. One generation of common garden was performed for replicate 3 of the control regime, to remove potential maternal effects. From these common gardens, 120 virgin females (dams) and 20 males (sires) were collected. Each sire was allowed to mate with six dams in a tomato leaf disk for 72 hours. This procedure was done in three consecutive days, in two successive weeks, in a total of six blocks. After mating, each dam was isolated on a tomato leaf disk to lay eggs for 72 hours. After 14 days, three dams were randomly selected per sire and four daughters per dam were transferred to leaf disks made from plants watered with 2mM of cadmium chloride. The experimental procedure used followed the protocol described above for quantifying spider mite performance.

### Statistical analyses

#### Spider mite performance

For the analyses of life-history traits (i.e., number of eggs laid, hatching rate and number of adult offspring) the average trait values of mites from each replicate population of the control selection regime (i.e., evolving on the cadmium-free environment) was determined for each environment (cadmium-free vs. 2mM of cadmium). Then, for the selection regimes evolving in the cadmium and heterogenous environmentsthe difference to the control was determined by subtracting from each measurement the average of the control population of the same replicate (for example: for a measurement of a trait on the cadmium-free environment from an individual of replicate 1 evolving on the heterogeneous environment it was subtracted the average trait value on the cadmium-free environment of the control population of replicate 1). These differences to the control were then analysed using general linear mixed models (GLMM), with the “glmmTMB” function of the “glmmTMB” package (Magnusson *et al*., 2017). Selection regime (i.e., evolving on the homogenous vs. heterogenous environment) and test environment (cadmium-free vs. 2mM of cadmium) were used as fixed factors, effective number of generations as covariate, leaf age and block as random factors and a gaussian error distribution. Models were simplified by removing non-significant terms and interactions. When there was a significant triple interaction (“selection regime” * “test environment” * “effective generation of selection”), we analysed the regimes evolving on the cadmium environment and those evolving on heterogenous environments separately (i.e., we divided the dataset by selection regime). Post-hoc comparisons to determine significant differences to control for each selection regime, on each test environment and at each generation were performed using the “emmeans” function of “lsmeans” package in R (Lenth & Lenth, 2016), comparing the slop of the difference between the control population and each selection regime to 0.

#### Habitat choice

The habitat choice was analysed using generalized linear mixed models (GLMM), with the “glmmTMB” function of the “glmmTMB” package (Magnusson *et al*., 2017). Selection regime was used as fixed factor, the effective number of generations as covariate, leaf age and block as random factors and a binomial error distribution, with choice of the cadmium-free environment coded as 0 and choice of the cadmium environment coded as 1. Because at generation 14 choice was only assessed after 24 hours and at generation 51 it was assessed after 24 and after 48 hours, we first analysed the differences in choice between 24 and 48 hours. Then, given the lack of significant differences (Chi^2^_1_= 0.04, P= 0.84), we only used the data obtained 24h after instalment for each generation for further analyses. Differences in the number of eggs laid on each available environment were analysed using similar models, with the response variable as the proportion of eggs laid per day on each environment (cadmium-free vs. 2mM of cadmium), using the “cbind” function in R.

#### Genetic variation

Using the full-sib half sib design allows to partition the phenotypic variance (VP) into additive (VA), dominance and maternal variance (VD+M) and environmental variance (VE) (Lynch & Walsh, 1998). Spider mites are haplodiploid organisms, which modifies variance partitioning as compared to diploid organisms (Liu & Smith, 2000; Magalhães et al., 2007): VA can be estimated as 2*σ^2^Sire (variance component of the sire) and VD+M can be estimated as 4*σ^2^Dam (Liu & Smith, 2000). Narrow sense heritability (h^2^) is calculated as 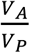 and broad sense heritability (which includes additive genetic variance and dominance and maternal genetic variance) as 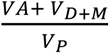.

Variance components and broad and narrow sense heritability were estimated using mixed models with the MCMCglmm package (Hadfield, 2010), using a poisson distribution. The full model included as random factors Sire, Dam nested within Sire and Block. A parameter expanded prior was used, since it is better for low variances (Gelman, 2006; Hadfield, 2010; de Villemereuil, 2018), with 500000 iterations of burnin, 750000 iterations and a thinning interval of 100, to decrease autocorrelation. All posterior traces were visually inspected to guarantee that the chains were mixing well. Credibility intervals were obtained using the function HPDinterval directly on the posterior samples after calculation of the observed heritability (de Villemereuil, 2018). To assess significance of the estimated variance components, comparisons between the deviance information criterion (DIC) of the full model with the DIC of models without Sire, Dam and the null model (fitting only the Block as random factor) was done (Henriques *et al*., 2021), since MCMCglmm constrains variance to be above 0. Models were considered significantly different if the DIC showed a difference above 2.

## Results

### Spider mite performance

#### Number of eggs

In the cadmium-free environment, populations evolving in the homogeneous environment laid a lower number of eggs than control populations (t.ratio= -6.13, P< 0.001; Fig. S2) and this was consistent between generations (Chi21= 0.01, P= 0.55). In contrast, in the same environment, the differences in number of eggs laid, between the populations evolving in heterogeneous environments and the control populations varied across generations (Chi21= 2.85, P= 0.04). Indeed, the number of eggs laid by the populations evolving in heterogeneous environments did not differ from that of the control populations at generation 12 (t.ratio= -1.53, P= 0.13; Fig. S1) but, it was lower than that of the control populations at generation 33 (t.ratio= -2.16, P= 0.03; Fig. S2). In the environment with cadmium, the number of eggs laid by populations evolving in the homogeneous environment was not significantly different from that of the control populations (t.ratio= -0.13; P= 0.90; Fig. S2), and this did not change with time as well (Chi21= 0.01, P= 0.55). The populations evolving in heterogeneous environments laid more eggs than control populations in the cadmium environment at generation 12 (t.ratio= 2.13; P= 0.03; Fig. S2) but not at generation 33 (t.ratio= 0.55; P= 0.58; Fig. S2).

#### Hatching rate

The hatching rate was not significantly different among selection regimes (Chi^2^_1_= 0.16, P= 0.69; Fig. S3), test environments (Chi^2^_1_ = 0.18, P= 0.67; Fig. S3), nor did it change from generation 12 to generation 33 (Chi^2^_1_= 2.03, P= 0.15; Fig. S3).

#### Number of adult offspring

Populations evolving in homogeneous environments and populations evolving in heterogeneous environments varied in their differences to the control populations across environments (interaction test environment * selection regime: Chi^2^_1_= 3.23, P= 0.07) and with time (interaction generation* selection regime: Chi^2^_1_= 3.35, P= 0.05). In the cadmium-free environment, both sets of populations produced a lower number of adult offspring than the control populations at generation 12 (t.ratio= -6.04, P< 0.001, t.ratio= 2.60; P= 0.01, for the populations evolving in the homogeneous and the heterogeneous environment, respectively; Fig. 1) and at generation 33 (t.ratio= -7.04, P< 0.001, t.ratio= -3.37; P= 0.001, for the populations evolving in the homogeneous and the heterogeneous environment respectively; Fig. 1). In the environment with cadmium, the number of adult offspring of populations evolving in the homogeneous environment was not significantly different, as compared to the control populations, at both time points (t.ratio= 0.86; P= 0.39, t.ratio= 1.38; P= 0.17; Fig. 1). In contrast, the populations evolving in heterogenous environments produced higher number of adult offspring than the control populations, but only at generation 12 (t.ratio= 2.94; P= 0.004 and t.ratio= 0.02; P= 0.98 for generations 12 and 33 respectively; Fig. 1).

**Figure 1.**
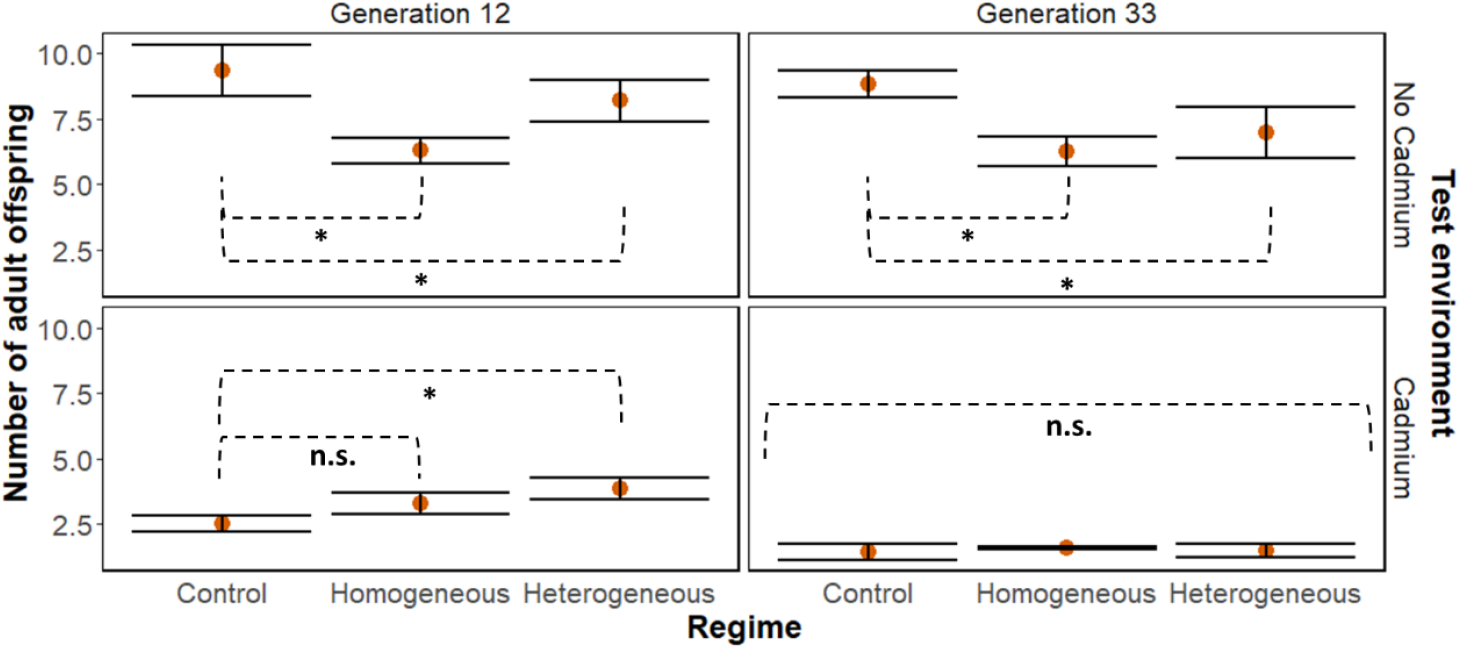
Adult offspring produced. Average number of adult offspring produced by *T. evansi* females from populations evolving in a cadmium-free environment (control), a homogeneous environment with cadmium and a heterogeneous environment (±se, N= 5 replicate populations), for 12 and 33 discrete generations (left and right panels, respectively), when tested in on plants with (top) or without (bottom) cadmium.

### Habitat choice

The number of females found in each test environment 24 hours the instalment did not differ significantly (Chi^2^_1_= 1.54, P= 0.21; Fig. 2). This result was also not affected by the selection regime (Chi^2^_2_= 0.67, P= 0.71; Fig. 2) nor the generation at which the experiment was done (Chi^2^_1_= 1.12, P= 0.29; Fig. 2). The number of eggs laid by *T. evansi* on the cadmium-free test environment was higher than that laid on the 2 mM cadmium test environment at generation 51 (Chi^2^_1_= 1.42, P< 0.001; Fig. S4), but not at generation 14 (Chi^2^_1_= 0.77, P= 0.13; Fig. S4), and, this pattern was similar across selection regimes (Chi^2^_2_= 0.66, P= 0.78 and Chi^2^_2_= 1.70, P= 0.43, for generation 14 and 51 respectively).

**Figure 2.**
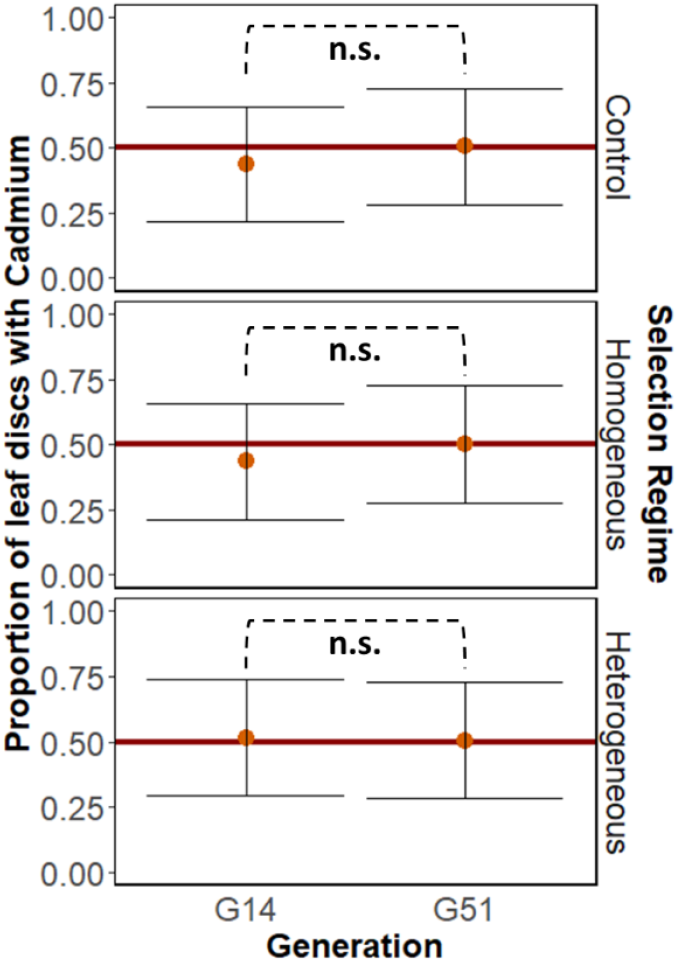
Habitat choice. Average proportion *T. evansi* females found on of leaf discs made from plants with 2 mM of cadmium or from cadmium-free plants 24h after being placed on the middle of a bridge connection them. Females were from populations evolving in a cadmium-free environment (control), on plants with cadmium (homogeneous) or on plant with and without cadmium (heterogeneous), (±se, N= 5 replicate populations), for 14 and 51 discrete generations (G14 and G51, respectively).

### Genetic variation

Both the number of eggs and number of offspring of the control regime in the cadmium environment had low heritability (Table 1). This result is further supported by the similarity of DICs when comparing models with and without the sire component (Table 2). Interestingly, models without the Dam component show a significant increase in the DIC (above 2), suggesting that the dominance and maternal variance have a high contribution to the phenotypic variance. Indeed, estimations for broad sense heritability are much higher than those for narrow sense heritability (Table 1). Nevertheless, it is important to note that the values are smaller than previously observed for these same traits (Magalhães *et al*., 2007).

**Table 1.** Mean heritability estimates (and corresponding credibility intervals) obtained from the posterior distribution. Note that MCMCglmm constrains the variance above 0, so significance cannot be obtained from comparing the overlap with 0.

| Trait | Additive genetic variance | Dominance and maternal variance | Narrow sense heritability | Broad sense heritability |
| --- | --- | --- | --- | --- |
| Number of eggs | 0.10971 | 0.39789 | 0.000118 (0.000001-0.000407) | 0.00183 (0.00066-0.0031) |
| Total offspring | 0.24384 | 0.62495 | 0.000125 (0.000001-0.000427) | 0.00141 (0.00044-0.00242) |

**Table 2.** DIC for different MCMC glmm models using either the complete model, removing the sire, removing the dam or removing both from the model.

| Trait | Full model | Without sire | Without dam | Without sire and dam |
| --- | --- | --- | --- | --- |
| Number of Eggs | 1889.349 | 1889.3 | 1909.669 | 1924.627 |
| Total Offspring | 1257.985 | 1258.074 | 1280.87 | 1302.638 |

## Discussion

Here used experimental evolution to test whether spider mites in either a homogenous or a heterogeneous environment adapt to tomato plants that have accumulated cadmium on their leaves. We find no changes in life-history traits or in habitat choice in response to cadmium accumulation in the spider mite *T. evansi* after 33 generations (although we find transient adaptation in the heterogeneous environment at generation 12). Additionally, evolving in environments with cadmium, being it homogenous or heterogeneous, entailed a cost in cadmium-free environments. Finally, we show that the life-history traits that we measured did not have significant heritability on cadmium.

Our results recapitulate that cadmium accumulation in tomato plants presents a strong selective pressure on *T. evansi* (Godinho *et al*., 2018, 2022b). Indeed, control populations have lower fecundity and lower production of adult offspring on environments with high cadmium, compared to cadmium-free environments. Despite this strong selective pressure, the performance of *T. evansi* on plants with cadmium did not increase after evolving 33 generations on such environment. This is in line with the absence of genetic variation for the measured life-history traits on cadmium in the control regime that we observe here. This regime was created from, and maintained in the same conditions (i.e., cadmium-free environment) as, the ancestral population that was created by merging 4 field populations using one-on-one crosses to ensure the representation of genotypes from all populations (Godinho *et al*., 2020). Moreover, we have documented intraspecific variability in the response to varying concentrations of cadmium accumulated on tomato plants in the populations used to create this population (Godinho *et al*., 2022a). Therefore, we were expecting high genetic variation for these traits in this population.

However, such absence of genetic variation for life-history traits in populations exposed to metal contamination has been observed in other species (Posthuma & Van Straalen, 1993). This absence of genetic variance may be due to the fact that metals are not part of the natural environment of organisms. Whereas this may be the case for most populations, there is also compelling evidence of organisms occurring in highly contaminated environments (Posthuma & Van Straalen, 1993; Boyd, 2009). Moreover, some studies show that organisms (Timmermans *et al*., 2005; Freeman *et al*., 2006; Fisker *et al*., 2011), including spider mites (Xu *et al*., 2021), adapt to metals. Also, we observed that the experimental populations were able to persist on environments with cadmium for many generations, producing enough offspring to be experimentally transferred to the following generation (i.e., backup populations were hardly used to maintain the density constant; Table S1, Fig. S1). Possibly, adaptation to cadmium involves different mechanisms and/or traits that we did not measure. Such, trait-specific adaptation patterns have been observed in other experimental evolution studies using spider mites (Magalhães *et al*., 2007; Marinosci *et al*., 2015). Furthermore, other studies revealing little evidence for phenotypic changes in life-history traits in response to cadmium did find a genetic signature of evolving in that environment (Ward & Robinson, 2005; Doria *et al*., 2022). These genetic changes were mostly associated with transporters involved in the excretion of metals and metallothionines, which are cadmium binding proteins implicated in its detoxification (Dallinger & Höckner, 2013; Doria *et al*., 2022). Other studies have shown that populations have adapted to selenium by accumulating a less toxic form of this metalloid (Freeman *et al*., 2006; Xu *et al*., 2021), but whether this detoxifying mechanism occurs in response to cadmium is still unknown. Even though some of the abovementioned studies do not concern the effect of metal accumulation on herbivores, we have previously shown that spider mite performance is best explained by the metal itself rather than by metal induced plant changes (Godinho *et al*., 2022a). It is thus reasonable to assume that the mechanisms may be similar. Therefore, incorporating genomic and transcriptomic analyses, as well as the measurement of other traits, in studies of herbivore adaptation to metal accumulation in plants will allow us to confirm whether these and other mechanisms contribute to the persistence of herbivore populations in metal polluted sites.

The lack of adaptation to cadmium in mites evolving in a heterogeneous environment composed of leaves with or without cadmium may be due to the reasons outlined above, but also to the potential migration load from leaves without cadmium to those with cadmium. Indeed, evolving in spatially heterogeneous environments with large differences in productivity between individuals on the different environments may hamper adaptation to the least productive environment due to excessive gene flow (Ronce & Kirkpatrick, 2001). In our set-up, spider mites perform much better in the environment without cadmium than in its presence, hence differences in productivity are high. Moreover, they could move freely between adjacent environments throughout their lifetime, hence we expect high levels of gene flow between leaves with and without cadmium. As such we expected adaptation to cadmium to be hampered in this environment, compared to populations evolving in a homogeneous environment.

Despite the lack of adaptation in the heterogeneous environment after 33 generations, we observed an increase in fecundity and in adult offspring production in the cadmium environment after 12 generations compared to the control selection regime, which was not the case for populations evolving in homogeneous environments. This suggests a transient response to cadmium accumulation in the populations evolving in heterogeneous environments. Other studies have documented such transient adaptation (Yona *et al*., 2012; Olazcuaga *et al*., 2022), including in spider mites adapting to heterogeneous environments with different host plant species (Bisschop *et al*., 2019). The occurrence of transient evolution in such environments could be due to the subsequent evolution of habitat choice, such that populations would progressively become less exposed to the stressful environment (Ravigné *et al*., 2009). However, habitat choice did not evolve in the spatially heterogenous environment, despite the potential clear benefits thereof, as observed in other studies in spider mites (Magalhães *et al*., 2009; Mortier & Bonte, 2020). Another explanation put forward for the occurrence of transient adaptation is that it allows population persistence in the early phases of colonization of a novel habitat, but then other, less costly mechanisms evolve later on that allow the establishment of the population in that environment. We have not been able to identify such a mechanism, meaning that whether this occurs in herbivores colonizing metal-accumulating plants remains to be confirmed.

Interestingly, despite no signs of adaptation, we observed a decrease in fecundity and in the production of adult offspring in the cadmium-free environment in populations evolving in environments with cadmium, either homogeneous or heterogeneous. This lends support to the hypothesis that other traits have evolved in the experimental selection regimes and these (unobserved) adaptations entailed a cost in the cadmium-free environment. For example, spider mites may have evolved a slower metabolism in environments with cadmium, improving its detoxification and increasing their life-spam. This slower metabolism would lead to a lower fecundity and adult offspring production across environments, even in cadmium-free environments. If so, evolving in environments with cadmium would reduce the growth rate of these herbivores, even when infesting non-accumulating plants, reducing herbivory overall.

Is adaptation to cadmium accumulation unlikely in spider mite populations? Bases on our study this question remains unanswered. Perhaps, in natural environments metal accumulating plants may serve as a refuge for spider mites when non-accumulating plants are temporally unavailable or overcrowded with competitors but long-term exposure to this contaminant leads to a dead end. Alternatively, adaptation may occur via unmeasured traits. This suggests that commonly measured life-history traits may not be sufficient to detect adaptation to metal accumulation and, thus, more and broader studies on this topic are needed.

## Acknowledgements

We would like to thank Cátia Eira, Maud Charlery and Inês Santos for the maintenance of the experimental populations, Lucie de Sousa for growing the plants and all the mite squad for fruitful discussions.

## Supplementary information

**Figure S1.**
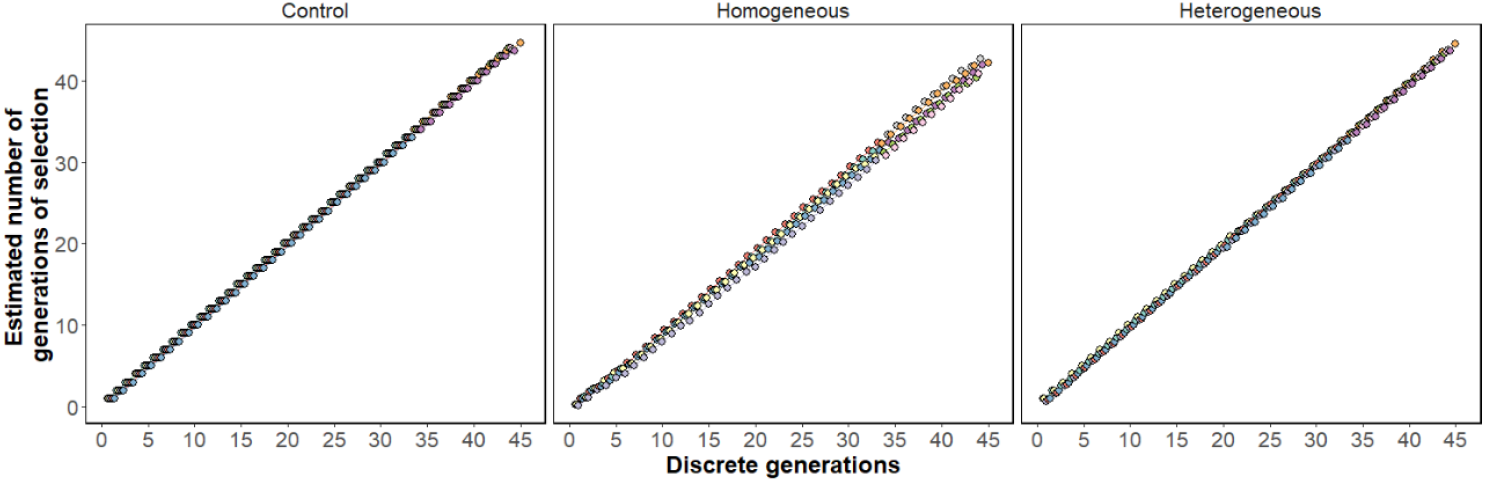
Effective number generations of selection thought time. Estimated number of generations of selection for each replicate of all selection regimes (control, homogeneous and heterogeneous) at each transfer (i.e., discreate generation).

**Figure S2.**
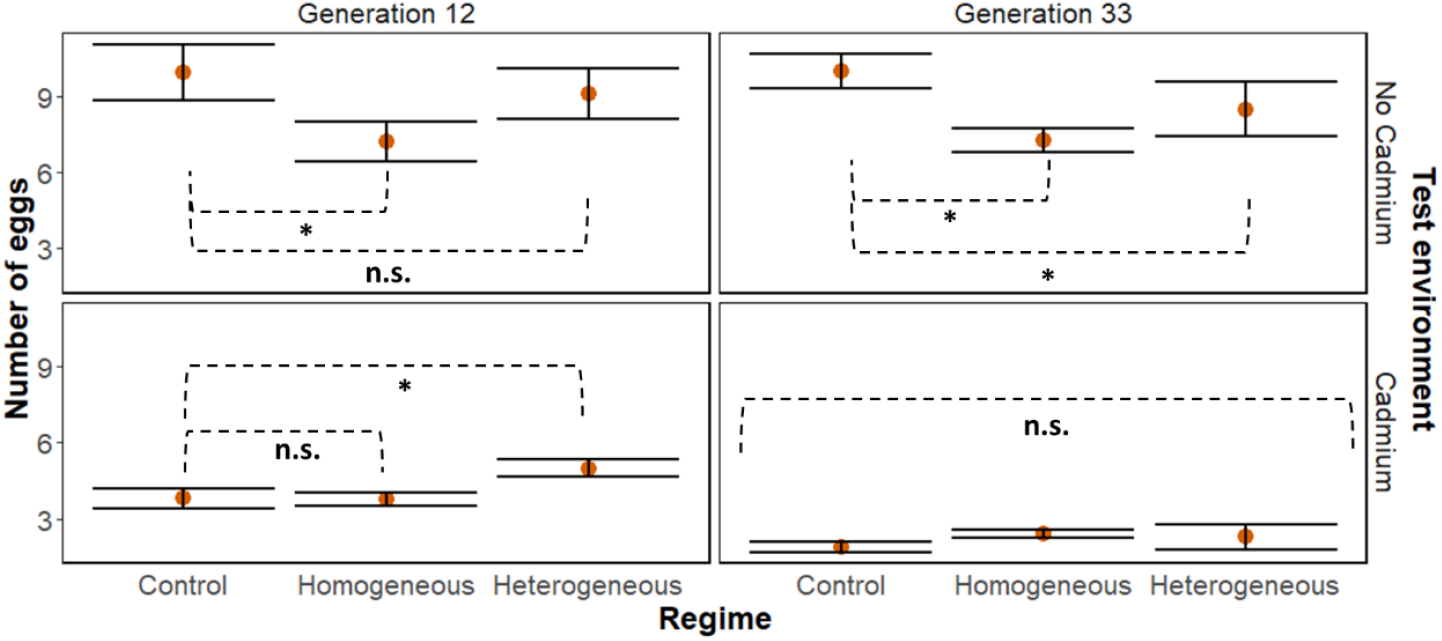
Number of eggs. Average number of eggs laid by *T. evansi* females from populations evolving in a cadmium-free environment (control), a homogeneous environment with cadmium and a heterogeneous environment (±se, N= 5 replicate populations), for 12 and 33 discrete generations, on plants with (top) or without (bottom) cadmium.

**Figure S3.**
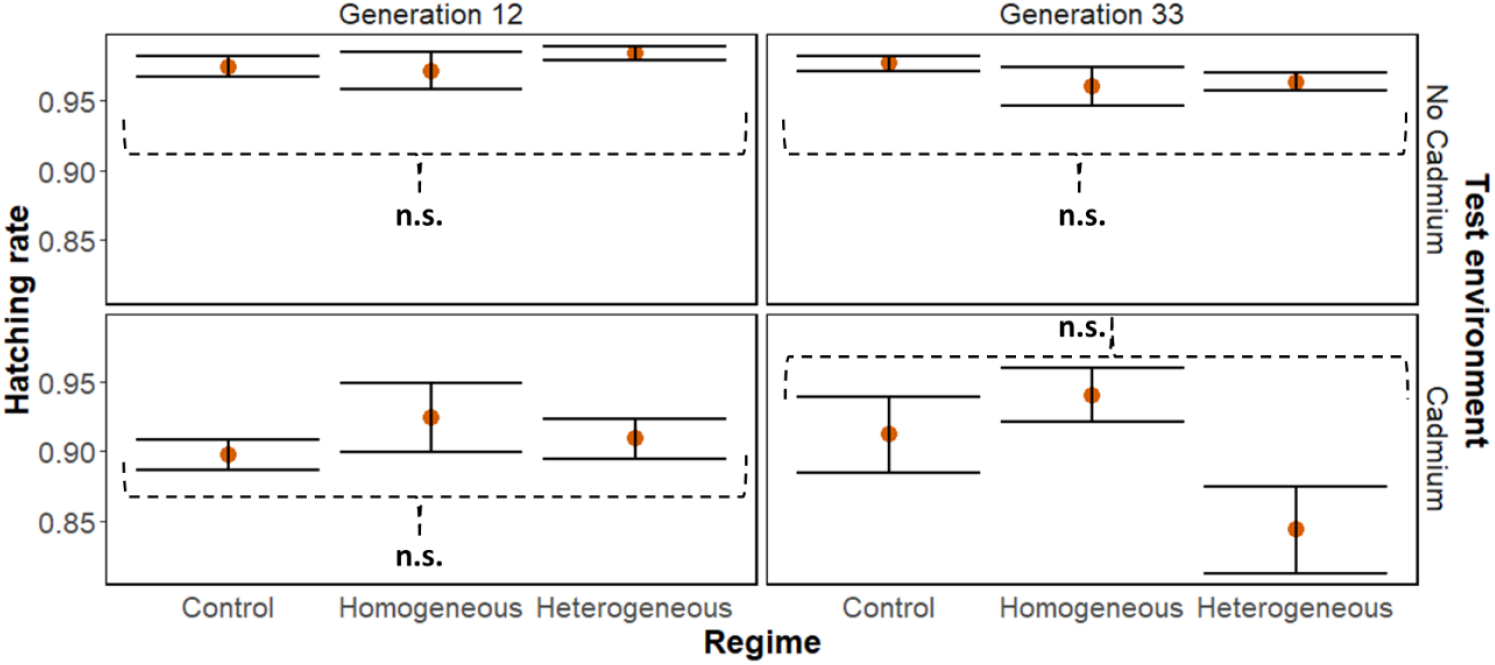
Hatching rate. Average proportion of eggs hatched for *T. evansi* females from populations evolving in a cadmium-free environment (control), a homogeneous environment with cadmium and a heterogeneous environment (±se, N= 5 replicate populations), for 12 and 33 discrete generations, on plants with (top) or without (bottom) cadmium.

**Figure S4.**
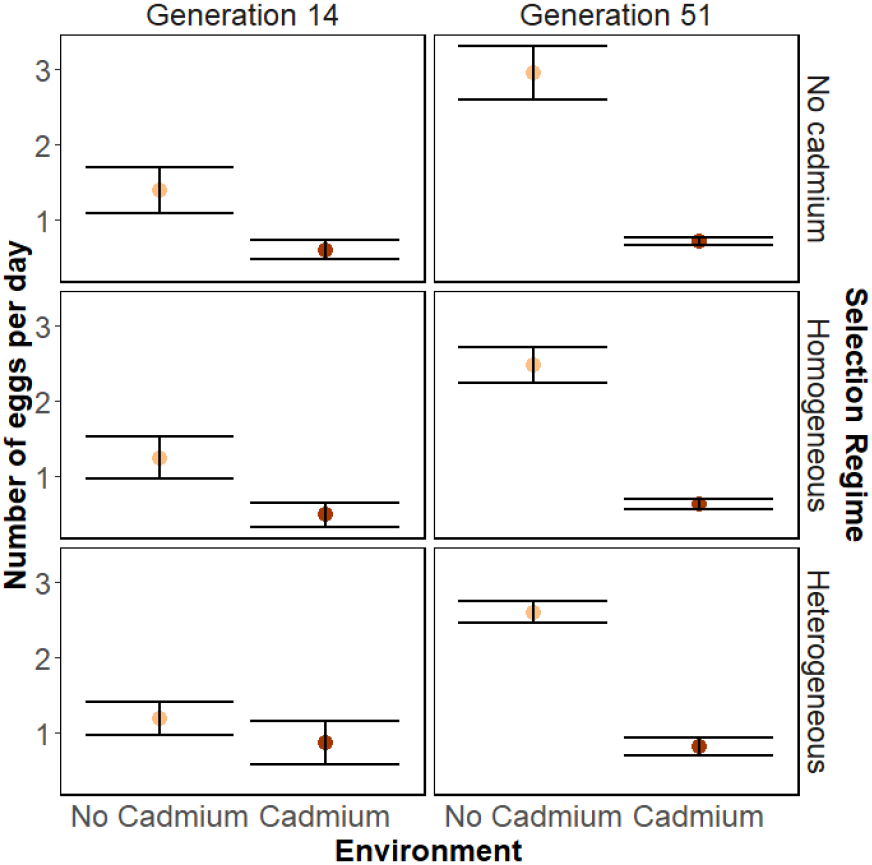
Choice of environment to lay eggs. Average number of eggs laid by *T. evansi* females from populations evolving in a cadmium-free environment, an environment with cadmium and a heterogeneous environment (±se, N= 5 replicate populations), on discs made from cadmium-free plants and on discs made from plants with 2 mM of cadmium, when given the choice between the two test environments, at generation 14 and 51.

**Table S1.** Effective number of generations of selection. Estimated number of generations of selection for each replicate population evolving on a cadmium-free environment (“Control”), an environment with plants supplied with 2 mM of cadmium (“Homogenous”) and a heterogeneous environment composed of leaves from cadmium-free plants and plants supplied with 2 mM of cadmium (“Heterogenous”) at the moment mated females were collected for the experiments. “Discrete generations”, correspond to the number to generations assuming no mites from the t-1 or the ancestral outbred population were reintroduced into the experimental evolution; “Effective number of generations of selection” corresponds to the number of generations elapsed accounting for the reintroduction of mites from the t-1 or the ancestral outbred population, calculated following Godinho et al. 2020.

|  | Control |  |  |  |  | Homogeneous |  |  |  |  | Heterogeneous |  |  |  |  |
| --- | --- | --- | --- | --- | --- | --- | --- | --- | --- | --- | --- | --- | --- | --- | --- |
|  | Replicate population |  |  |  |  | Replicate population |  |  |  |  | Replicate population |  |  |  |  |
|  | 1 | 2 | 3 | 4 | 5 | 1 | 2 | 3 | 4 | 5 | 1 | 2 | 3 | 4 | 5 |
| Discrete generation | Effective number of generations of selection |  |  |  |  |  |  |  |  |  |  |  |  |  |  |
| 12 | 12 | 12 | 12 | 12 | 12 | 10,31 | 10,33 | 9,94 | 11,40 | 11,15 | 11,46 | 12 | 11,62 | 11,77 | 12 |
| 14 | 14 | 14 | 14 | 14 | 14 | 12,31 | 12,33 | 11,56 | 13,40 | 13,15 | 13,46 | 14 | 13,62 | 13,77 | 14 |
| 33 | 33 | 33 | 33 | 33 | 33 | 31,31 | 30,66 | 30,14 | 32,40 | 31,35 | 32,46 | 32,57 | 32,46 | 32,77 | 32,60 |
| 51 | 50,63 | 51 | 51 | 50,73 | 50,60 | 47,55 | 47,13 | 47,81 | 49,63 | 48,91 | 50,46 | 50,30 | 50,46 | 50,77 | 50,60 |

